# Untangling the complexity of priority effects in multispecies communities

**DOI:** 10.1101/2021.03.29.437541

**Authors:** Chuliang Song, Tadashi Fukami, Serguei Saavedra

## Abstract

The history of species immigration can dictate how species interact in local communities, thereby causing historical contingency in community assembly. Since immigration history is rarely known, these historical influences, or priority effects, pose a major challenge in predicting community assembly. Here, we provide a graph-based, non-parametric, theoretical framework for understanding the predictability of community assembly as affected by priority effects. To develop this framework, we first show that the diversity of possible priority effects increases super-exponentially with the number of species. We then point out that, despite this diversity, the consequences of priority effects for multispecies communities can be classified into four basic types, each of which reduces community predictability: alternative stable states, alternative transient paths, compositional cycles, and the lack of escapes from compositional cycles to stable states. Using a neural network, we show that this classification of priority effects enables accurate explanation of community predictability, particularly when each species immigrates repeatedly. We also demonstrate the empirical utility of our theoretical framework by applying it to two experimentally derived assembly graphs of algal and ciliate communities. Based on these analyses, we discuss how the framework proposed here can help guide experimental investigation of the predictability of history-dependent community assembly.

## Introduction

An ecological community’s assembly history is defined by the order and timing of species arrival (Fukami, 2015). Arrival timing and order have a strong random component (Gould, 1990; Sprockett *et al.*, 2018), making it difficult to predict which assembly history will take place. The large number of conceivable assembly histories has highlighted the necessity of understanding how history affects the diversity, composition, and functioning of an ecological community (Fukami & Morin, 2003; Toju *et al.*, 2018; Carlström *et al.*, 2019; Clay *et al.*, 2020). In this vein, mounting data suggests that assembly history affects which species survive because species can interact in local communities differently depending on arrival order and timing (Fukami, 2015), the phenomenon known as *priority effects*. Much of our present understanding of priority effects, however, is based on the simple 2-species Lotka-Volterra model and its variants (Gilpin & Case, 1976; May, 1977; Mordecai, 2013; Fukami *et al.*, 2016; Wittmann & Fukami, 2018; Ke & Letten, 2018; Song *et al.*, 2020; Ke & Wan, 2020). It remains unclear how the diversity and complexity of priority effects expand beyond these models and when considering multispecies communities (Lawton, 1999; Fukami, 2015).

For example, arrival order dictates one of the three well-studied outcomes of 2-species competition in the Lotka-Volterra model (Figure 1A) (Case, 2000): (I) coexistence (two species coexist regardless of who arrives first); (II) deterministic exclusion (the same species always excludes the other regardless of who arrives first); and (III) history-dependent exclusion (the species that arrives first excludes the other). Though not as widely recognized, three other outcomes are also possible in 2-species competition: (IV) two species eventually coexist despite requiring a specific arrival order; (V) only one species survives or two species coexist depending on who arrives first; and (VI) the species that arrives late always replaces the species that arrives early. The presence of these other possibilities has been supported by empirical evidence (Drake, 1991; Warren *et al.*, 2003; Amor *et al.*, 2020; Angulo *et al.*, 2020). Of these six scenarios, (III), (V) and (VI) all represent priority effects. The common omission of cases (V) and (VI) in theoretical studies is a consequence of assuming that model parameters are fixed and history-independent (Rudolf, 2019; Zou & Rudolf, 2020). In reality, interaction strengths can be history-dependent (Rasmussen *et al.*, 2014; Poulos & McCormick, 2014; Vannette & Fukami, 2017; Carter & Rudolf, 2019; Sniegula *et al.*, 2019). Thus, the scope of priority effects is largely underestimated by traditional parametric models assuming history-independent interaction strengths.

**Figure 1:**
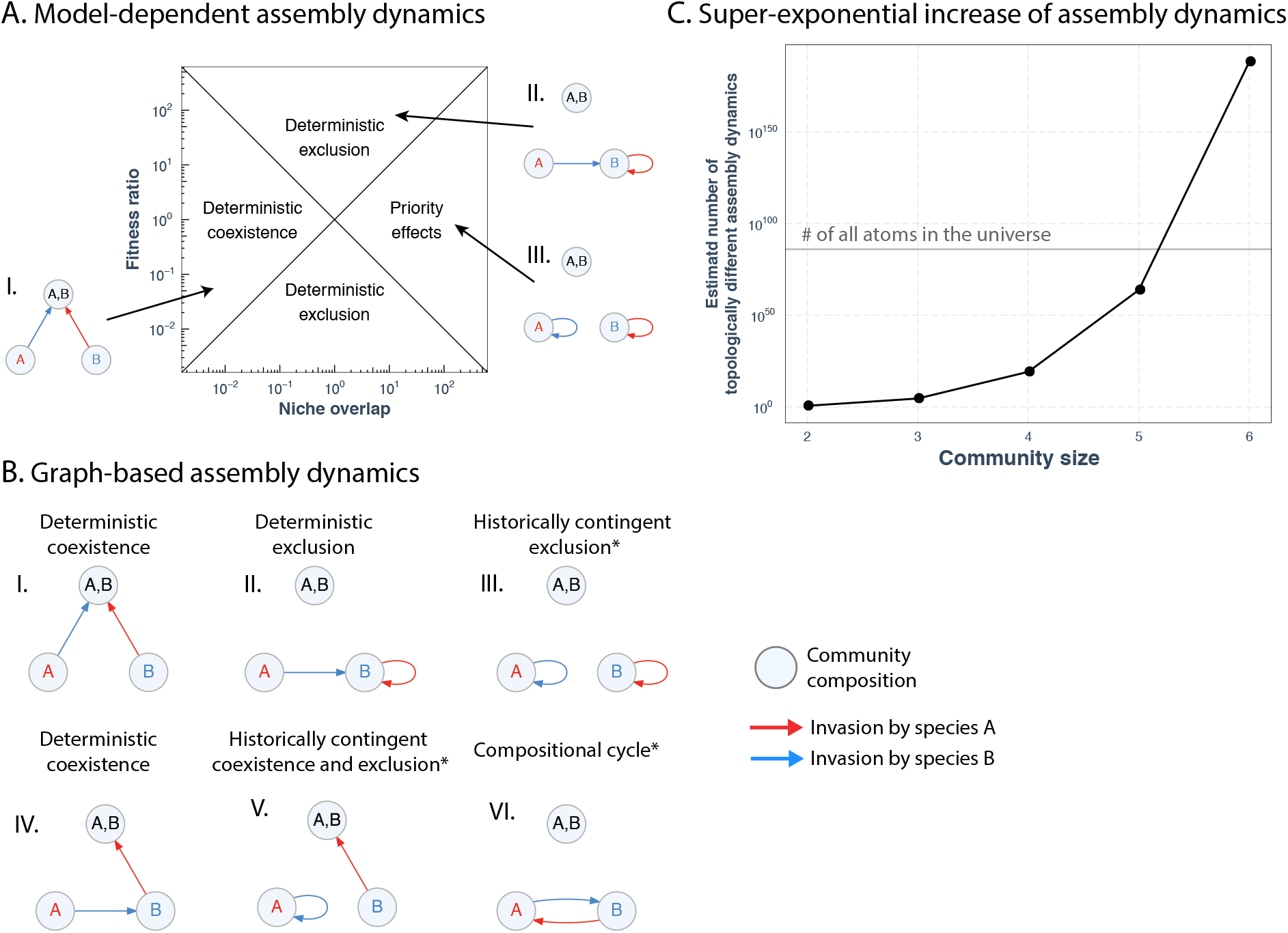
Non-parametric graph-based approach. Panel (**A**) presents the traditional model-based phase diagram of assembly dynamics in 2-species communities using coexistence theory. The balance between niche overlap (*x* axis) and fitness ratio (*y* axis) determines the types of assembly dynamics: (I) deterministic existence, (II) deterministic exclusion, or (III) priority effects. Panel (**B**) presents all six types of possible assembly dynamics (model-free) using the graph-based approach. In an assembly graph, the nodes represent the species combinations, while the links represent how the species combination change into another combination after a non-resident species invades. Take Case (II) for example: if species *B* (blue line) invades the community with species *A* (red circle), species *A* is excluded and the community composition changes into species *B* (hence the arrow goes from red circle to blue circle); and if species *A* (red line) invades the community with species *B* (blue circle), species *A* cannot establish and the community composition remains the same (hence the arrow turns back on itself). Cases (III), (V), and (VI) are all priority effects (marked with * symbol) as at least two different assembly histories can lead to different compositions. Panel (C) plots how the diversity of priority effects increases with larger community size. We find a super-exponential increase of diversity of topologically different assembly dynamics (Eqn. 1).

Moreover, the scope of priority effects quickly increases as more species are considered. For example, even with only three species, 41,979 outcomes are theoretically possible, the bulk of which constitute priority effects (Figure 1D). In general, as the number of species increases, the number of alternative outcomes increases super-exponentially (Figure 1D). The core reason behind this super-exponential increase is that, while all history-independent dynamics are alike, priority effects can be unique in their own way (Fukami, 2015). Only when all conceivable assembly histories result in the same community composition does history-independent dynamics arise. The extremely large number of assembly dynamics in multispecies communities makes the prospect of predicting community assembly daunting (Lawton, 1999; Srivastava, 2018).

While knowing the exact assembly dynamics operating in a community may be an impossible task indeed, we may still be able to understand the predictability of community assembly affected by priority effects: the extent to which we can predict community composition given the stochasticity of assembly history. For instance, if in one case all different assembly histories lead to different outcomes, and if in another case only one assembly history leads to a different outcome, the latter case has a much more predictable outcome than the former (Margalef, 1973). Understanding the predictability of community assembly is not only of basic interest to ecologists, but can also aid applications of community ecology for ecosystem management, including ecological restoration, biological control, and the medical treatment of the gut microbiome (Song & Saavedra, 2018; Sprockett *et al.*, 2018; Rohr *et al.*, 2020; Deng *et al.*, 2021). Yet, a theoretical framework is largely lacking for quantifying the predictability of community assembly in this context.

Here, we introduce a non-parametric graph-based approach to studying the diversity of priority effects in multispecies communities. Taking this approach, we propose an information-based metric to quantify community predictability, with a focus on the influence of priority effects. We show that the predictability of community assembly has two types of regularities: (1) The higher the invasion times, the higher the predictability; (2) Predictability of final composition and of temporary changes in community compositions are strongly correlated. We then show that the consequences of priority effects for community predictability can be classified into four basic sources: the number of alternative stable states, the number of alternative transient paths, the length of compositional cycles, and the presence or absence of an escape from cycles to stable states. We demonstrate both theoretically and empirically that this classification allows for accurate explanation of community predictability. Finally, we discuss how our results can be used to guide experimental studies for community assembly.

### A non-parametric graph-based approach

To capture the full diversity of priority effects, we introduce a non-parametric graph-based approach. This approach maps any assembly dynamics uniquely onto an assembly graph, where nodes represent combinations of coexisting species and directed links represent how species combinations change when a new species invades. This form of graph presentation is not new (Hang-Kwang & Pimm, 1993), but has been largely underused. We illustrate this approach with two hypothetical species, where species are denoted as *A* and *B* (Fig. 1B). This illustrative assembly graph has 2^2^ − 1 = 3 nodes: {*A*}, {*B*}, and {*A*, *B*}, which represent all possible species combinations. The link starting from community (node) {*A*} is generated by the invasion of species *B*, which can lead to all three possible species combinations (similarly for community {*B*}). In turn, community {*A*, *B*} does not have any directed link since all species are present in this community. For example, case (II) in Figure 1B shows the graph representation of deterministic exclusion: invasion from species *B* into community {*A*} leads to community {*B*}, whereas invasion from species *A* into community {*B*} also leads to community {*B*}. Note that there are in total 3 × 3 = 9 assembly dynamics without considering which species is named as {*A*} or {*B*} (which is arbitrary). Formally, this only considers *topologically unique* assembly dynamics (graphs), where an assembly graph is unique up to the ordering of species labels. Thus, with 2 species, there are only 6 topologically unique assembly dynamics (Fig. 1B).

We now generalize our approach to multispecies communities. For simplicity, we present this extension with 3 species with species denoted as *A*, *B*, and *C*. The assembly graph has 2^3^ − 1 = 8 nodes, representing all possible species combinations. In a community with a single species ({*A*}, {*B*}, or {*C*}), there are two outgoing links representing invasions by the other two species, respectively. For instance, in community {*A*}, one of the two outgoing links represents the invasion by species *B*, which can lead to all three possible combinations of species *A* and *B* ({*A*}, {*B*}, or {*A,B*}). Then, in a community with two species ({*A,B*}, {*A,C*}, or {*B,C*}), there is one outgoing link representing the invasion by the only species not present in the community. That is, in community {*A,B*}, the only outgoing link represents the invasion by species *C*, which can lead to all eight possible combinations of species *A*, *B*, and *C*. Note that community size may decrease after an invasion. Finally, in the community with all three species ({*A,B,C*}), there is no outgoing link since all species are present. In total, there are 41,979 topologically different assembly dynamics for just 3 species.

The same procedure above applies to an arbitrary number of species. The assembly graph for *S* species has (2^*S*^ − 1) nodes. The nodes representing communities with *n* species have (*S* − *n*) outgoing links. Each of the associated outgoing links from the node with *n* species can possibly lead to (2^(*n*+1)^ − 1) nodes. Then, out of the (2(*n*+1) − 1) nodes, only one node contains (*n* + 1) species, 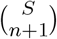 nodes contain *n* species, while all the other nodes contain less than *n* species. As a first-order of approximation, the diversity (number) of topologically different assembly dynamics can be calculated as (see Appendix A for derivation):

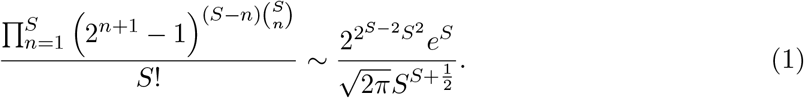

Figure 1D shows how the diversity of priority effects scales with the number of species. Note that already with 6 species, the diversity is significantly greater than the total number of atoms in the entire universe (Kragh, 2003).

### Priority effects and the predictability of community assembly

Priority effects give rise to assembly dynamics that are not fully predictable. To quantify this lack of predictability owing to priority effects, it is necessary to define both the pool of possible assembly histories for a given community and the type of uncertainty to analyze (Fig. 2). Focusing on the pool of assembly histories, if an infinite number of invasions is possible, the pool of assembly histories is also infinite. While this assumption is typically applied to allow statistical convergence, it is a rather strong assumption that is often not met (Hubbell, 1997; Capitán *et al.*, 2009; Serván & Allesina, 2020). As the assembly graph fully determines the trajectory of community composition given any assembly history, we can study any arbitrary pool of assembly histories (Appendix B). Here, for simplicity, we assume that each species invades *m* times. With 2 species, if species can only invade *m* = 1 time, the pool of possible assembly histories consists of only two assembly histories: 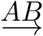 (meaning species *A* invades first and then species *B* invades) and 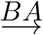 (Fig. 2B). But if we have 3 species that each invade *m* = 2 times, the pool of possible assembly histories consists of 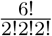 = 90 assembly histories. For example, 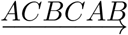 is a possible assembly history, meaning that species *A* invades first, then species *C* invades, and so on until species *B* invades last. By changing the number of invasion attempts (*m*), it is also possible to answer how large *m* has to be to effectively generate the same effects as infinite invasions. Similarly, our framework can be easily adapted to incorporate more ecological complexity, such as that some species arrive with higher frequency than others (see Appendix B for details).

**Figure 2:**
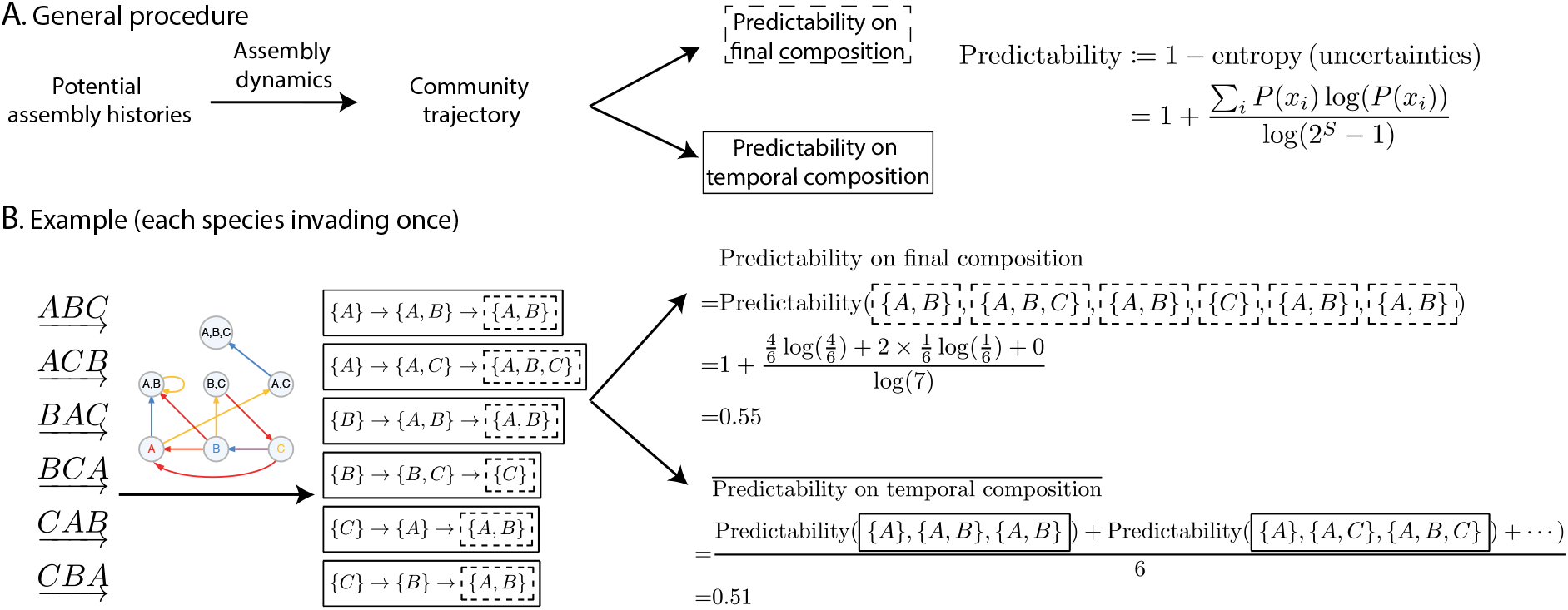
Quantifying the predictability of community assembly. Panel (**A**) presents the general procedure used for the calculation: First, based on a given uncertainty (either final composition or temporary compositions), we define a pool of potential assembly histories. Second, we need to know the assembly dynamics of the community. Third, each assembly history produces its community trajectory (how community composition changes with an invading species) based on the assembly dynamics. Last, we compute predictability on the final composition across the community trajectory (boxed with dashed lines) and on the temporary compositions along the community trajectories (boxed with solid lines). The definition of predictability is based on normalized entropy (see the mathematical definition in Eqn. 2 or in the figure). Panel (**B**) illustrates this procedure with an example of a 3-species community. First, we define the pool of potential assembly histories as each species invades only once. This gives us six potential assembly histories: 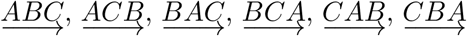 Second, we choose a 3-species assembly dynamics for illustration. Third, we show the community trajectories defined by assembly history and assembly dynamics. For example, the community trajectory is {A}→ {A, B} → {A, B} for the assembly history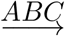. Last, we calculate the community predictability: the predictability on final composition isthepredictability of the states ({*C*}, 4{*A, B*}, {*A, B, C*})–the final compositions in each community trajectory (boxed in dashed lines), which is 0.55; and the predictability on temporal composition is the average predictability of six trajectories corresponding to six assembly histories (boxed in solid lines), which is 0.51.

Shifting our focus to the type of uncertainty to analyze, we suggest that, given a pool of assembly histories, it is possible to measure two types of uncertainties related to community composition. The first type is the uncertainty associated with the final community composition across all assembly histories. The second type is the uncertainty associated with transient community compositions along all assembly histories. Here, to quantify the predictability of assembly dynamics, we adopt a normalized information entropy metric (Rohr *et al.*, 2016). Although there are many alternative uncertainty metrics (Vellend, 2016), information entropy has been useful to quantify and explain different ecological processes (O’Connor *et al.*, 2019; Marleau *et al.*, 2020; Zu *et al.*, 2020; Margalef, 1973). We define the predictability of an assembly dynamics as

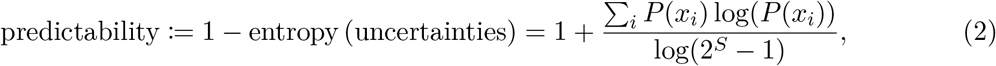

where *x_i_* is a species combination and *P* (*x_i_*) is the probability that combination *x_i_* occurs (Fig. 2A). The entropy is normalized to [0, 1] to ensure interpretability across different community sizes. Thus, a predictability of one (resp. zero) implies that that there is no (resp. full) uncertainty about the assembly dynamics.

To illustrate this measure, let us consider the assembly dynamics defined by case (IV) shown in Figure 1B, where each species invades only once. In this example (Fig. 2B), if the assembly history 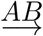 takes place, the trajectory of the community is from {*A*} → {*B*}. Instead, if assembly history 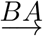 takes place, the trajectory is from {*B*} → {*A, B*}. Therefore, the predictability of the final community composition is given by predictability 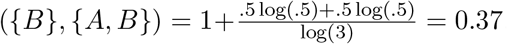. Similarly, the predictability of the assembly trajectory (the temporary community compositions) are predictability({*A*}, {*B*}) = .37 and predictability({*B*}, {*A, B*}) = .37. Figure 2B illustrates the process of quantifying this predictability for a set of 3 species. The agreement between the two types of predictability is not a coincidence. Figure 3A shows the strong correlation between the two types of predictability across all possible assembly dynamics for 3 species. The level of correlation is invariant across different pools of potential assembly histories. Thus, without loss of generality, hereafter our results are based on the first type of predictability (i.e., on the final community composition). Communities become more predictable when species are allowed to invade multiple times. Figure 3B shows how the distribution of predictability across all possible assembly dynamics with 3 species increases as a function of the number of invasion attempts.

**Figure 3:**
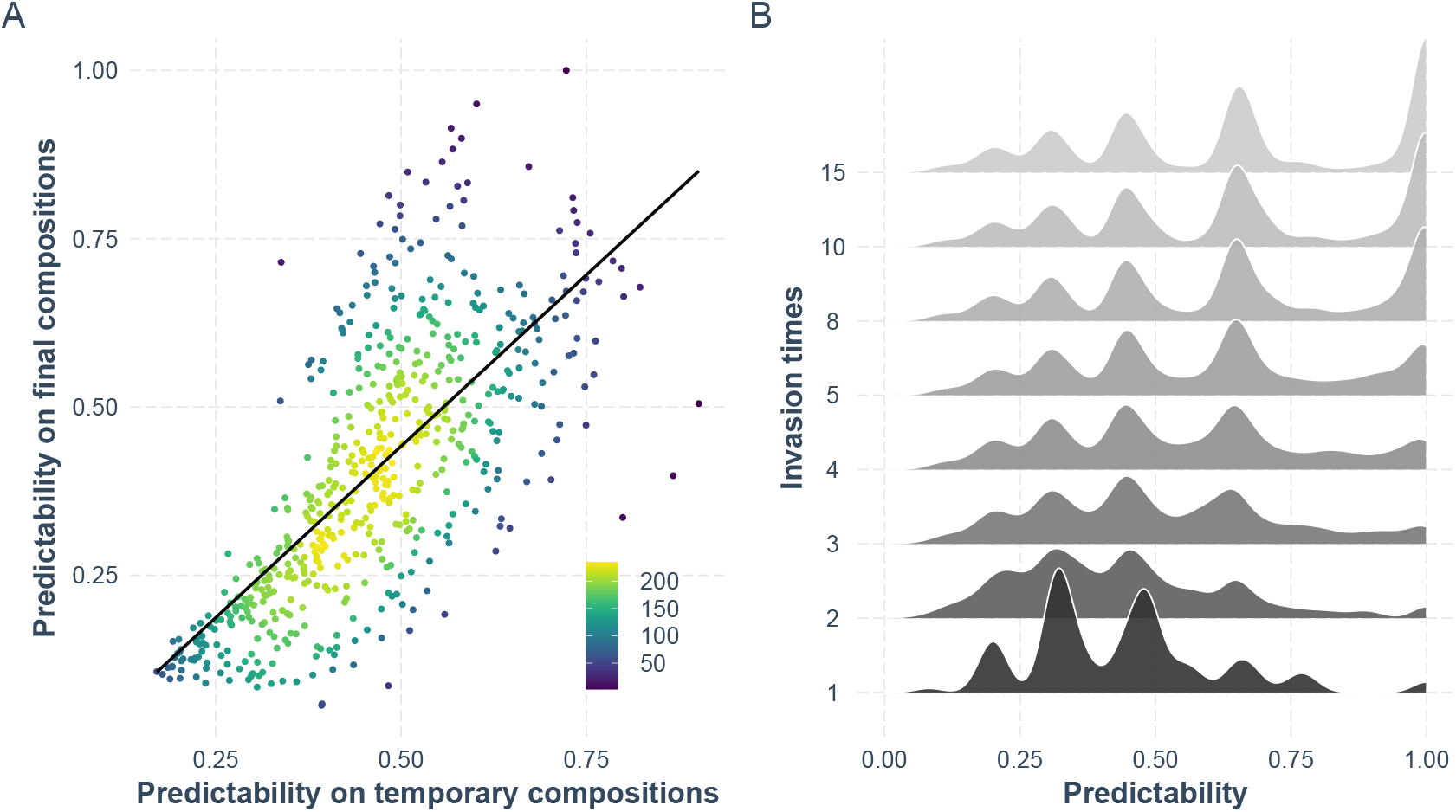
Regularities in the predictability of community assembly. Panel (**A**) shows the correlation (Pearson correlation = .72 with 95% confidence interval [.68, .75]) between the predictability of community assembly on the temporary compositions (*x* axis) and on the final compositions (*y* axis). Recall that all assembly dynamics with predictability less than 1 correspond to priority effects. The color map represents the number of data points. Panel (**B**) plots the predictability distribution of assembly dynamics as a function of invasion times. The predictability (*x* axis) corresponds to the predictability on the final composition. The distribution converges to a stationary distribution with increasing invasion times, i.e., the multiple humps in the distributions would be fixed.

### Classifying priority effects

As shown above, there are significant differences in the predictability of community assembly. Here, we show that we can decompose community predictability into four basic sources: the number of alternative stable states, the number of alternative transient paths, the length of compositional cycles, and the presence or absence of an escape from cycles to stable states.

Alternative stable states occur when a community has more than one stable composition (left column of Figure 4A) (Gilpin & Case, 1976; Schröder *et al.*, 2005; Schooler *et al.*, 2011). Alternative transient states occur when there are more than one assembly histories (or trajectories) from the founding species to the stable states (middle column of Fig. 4A) (Fukami & Nakajima, 2011, 2013; Sarneel *et al.*, 2019). Compositional cycles occur when the assembly histories involve cyclic sequences of community composition (right column of Fig. 4A) (Schreiber & Rittenhouse, 2004; Fox, 2008). Our graph-based approach can identify these dynamical sources as topological features. That is, alternative stable states arise when the assembly graph has more than one sink (nodes that have incoming links but no outgoing link); alternative transient states arise when more than one directed path exist from single species to a sink; while compositional cycles occur when directed cycles exist in the assembly graph. These three dynamical sources have already been hypothesized to be major drivers of priority effects (Fukami, 2015).

**Figure 4:**
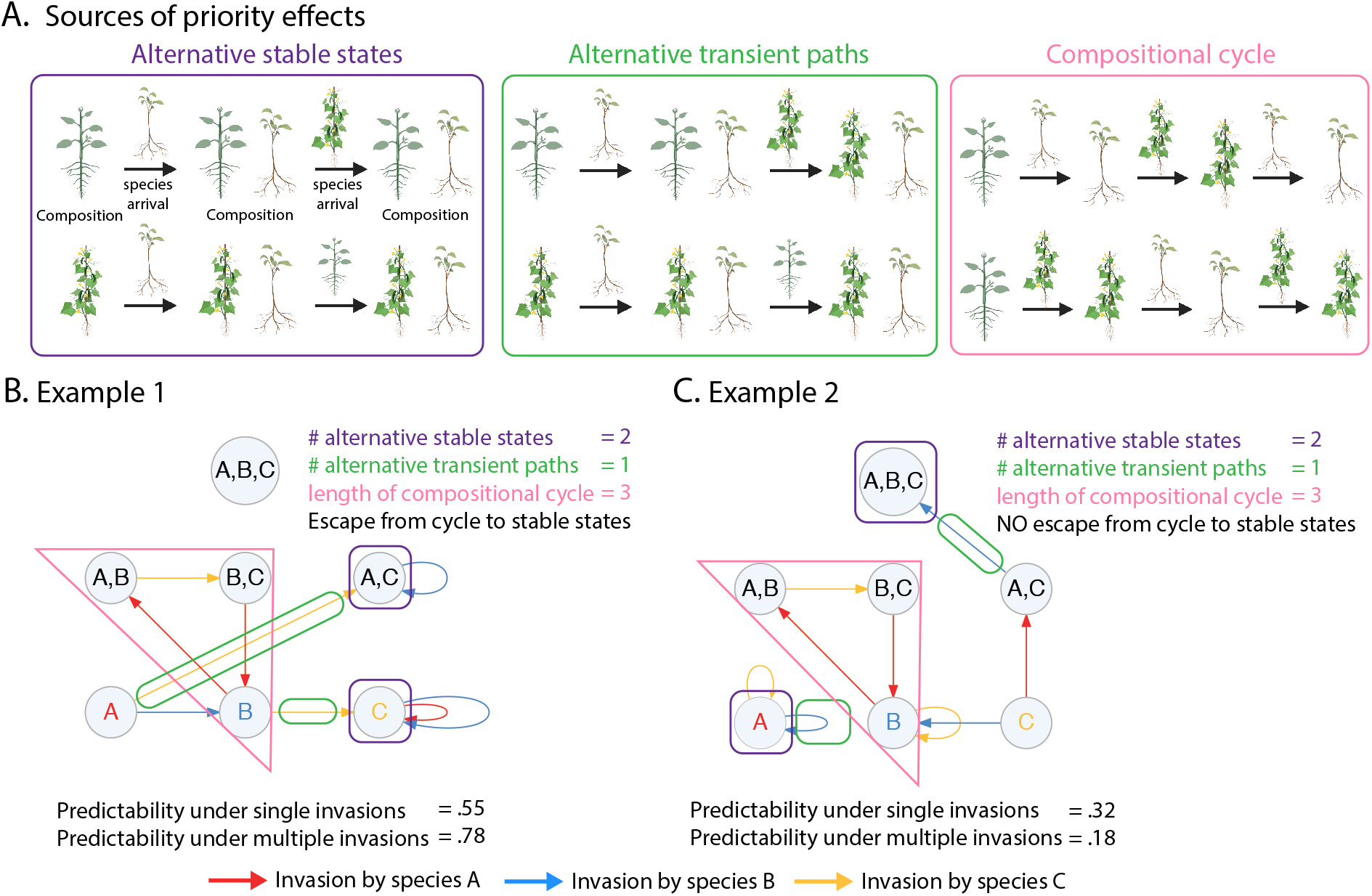
Dynamical sources of priority effects. Panel (**A**) presents the three known dynamical sources of priority effects (Fukami, 2015): alternative stable states, alternative transient paths, and compositional cycles. We illustrate these sources with two plants (cartoons). Panels (**B**) and (**C**) illustrate how these dynamical sources can be represented as topological features of assembly graphs. Panel (**B**) shows an assembly graph for 3 hypothetical species. It has two alternative stable states ({*A, C*} and {*C*}; boxed in purple lines), one transient path for each stable state ({*A*} → {*A, C*} and {*A*} → {*B*} → {*A, C*}; boxed in green lines), and one compositional cycle of length 3 ({*B*} → {*A, B*} → {*B, C*} → {*B*}; boxed in purple lies). Panel (**C**) shows an assembly graph for another set of 3 hypothetical species (**B**). It also has two alternative stable states ({*A*} and {*A, B, C*}; boxed in purple lines), one transient path for each stable state ({*A*} → {*A*} and {*C*} → {*A, C*} → {*A, B, C*}; boxed in green lines), and one compositional cycle of length 3 ({*B*} → {*A, B*} → {*B, C*} → {*B*}; boxed in purple lies). However, the assembly graphs in Panels (**B**) and (**C**) have drastically different predictability. The cause of this difference is a missing fourth topological feature: escape from compositional cycle to stable states. In Panel (**B**), the system can escape from the compositional cycles to stable states (i.e., from {*B*} to {*C*}). In Panel (**C**), the system cannot escape.

The three dynamical sources are not mutually exclusive, implying that assembly dynamics can potentially exhibit all of these sources. For example, the assembly graph for the 3 species shown in Figure 4B has two alternative stable states ({*A, C*} and {*C*}), one transient path for each stable state ({*A*} → {*A, C*} and {*A*} → {*B*} → {*A, C*}), and one compositional cycle of length 3 ({*B*} → {*A, B*} → {*B, C*} → {*B*}). However, the picture is not complete with only these three sources. Figure 4C shows another example of an assembly graph for 3 species with the same types and number of dynamical sources as shown in Figure 4B. Nevertheless, the assembly dynamics in Figure 4C has lower predictability than the one shown in Figure 4B. The difference between these two cases is given by the fact that only in Figure 4B, the assembly dynamics exhibits the possibility to escape from a cycle to a stable state (i.e., the trajectory can escape the compositional cycle ({*B*} → {*A, B*} → {*B, C*} → {*B*}) into a stable state ({*C*}). Because cycles are less predictable than stable states in general, this possibility to escape can increase the predictability of assembly dynamics.

Therefore, to classify priority effects by their predictability, we propose to use the four dynamical sources (topological features): alternative stable states, alternative transient states, compositional cycles, and the presence or absence of an escape from cycles to stable states. We used a neural network to carry out this classification. In short, the architecture of the neural network is as follows: the input layer is a four-dimensional vector, which encodes the four topological features of an assembly graph; the five hidden layers all have ReLU activation; and the output layer is the explained predictability. Appendix C provides a more detailed description of the neural network. For simplicity, we measure the explanatory (classification) power using the the correlation between observed and classified predictability in the out-of-sample test set. Focusing on the case of 3 species, we found that the classification works better for assembly dynamics with multiple invasions. Specifically, the classification displayed an explanatory power of 0.57 and 0.98 for single (*m* = 1; Fig 5A) and multiple (*m* = 15; Fig 5B) invasions, respectively. The explanatory power increases with the number of invasions and reaches a plateau around *m* = 8 invasions (Figure 5C). These results are qualitatively the same for larger communities (Appendix D).

**Figure 5:**
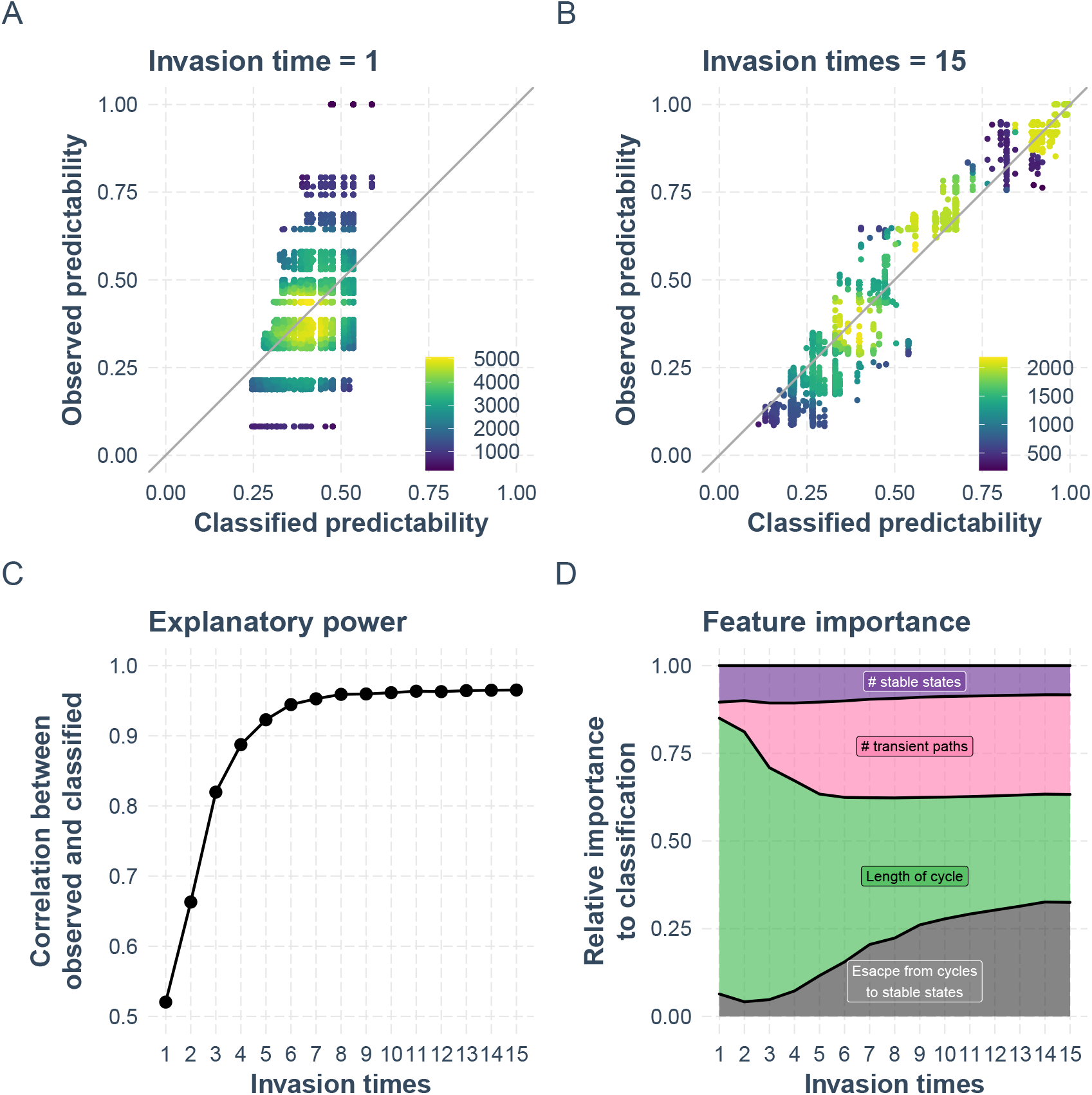
Explaining the predictability of community assembly using topological features. This figure illustrates the out-of-sample explanation power of the four dynamical sources (topological features) in classifying the predictability on the final composition for 3 species. We used 80% out of 41,979 assembly dynamics as the training set, while the remaining 20% was used as the test set (the classification). We used a neural network to achieve the maximum classification power (see Appendix C for details). Panel (**A**) shows the explanation power for the pool of assembly histories where each species invades only once. The *x* axis represents the classified predictability (from the test set) using the four topological features, while the *y* axis represents the observed predictability. The color of the point indicates how many assembly graphs are represented by the point. The gray line is the 45-degree line, representing that the observed predictability equals the classified predictability. Panel (**B**) shows the explanation power for the pool of assembly histories where each species invades 15 times. Panel (**C**) shows how the explanatory power increases with invasion times and reaches a plateau fast. We indicate the explanatory power using the correlation between the observed and classified predictability (using different pools of potential assembly histories). Panel (**D**) shows the relative importance of each topological feature (dynamical source) to the explanation power.

Finally, we used a regression-based scheme (Grömping *et al.*, 2006) to quantify the relative importance (contribution) of each topological feature (dynamical source) to the classification power of priority effects. Figure 5D shows that the relative importance of topological features changes with the number of species invasions. Additionally, this figure shows that the relative importance of the number of stable states remains constant across invasion times, while the relative importance increases for both the number of transient paths and the presence of an escape from cycles to stable states, but it decreases for the length of cycles. Moreover, this result shows that the relative importance of the number of stable states can be smaller than the combined importance of the other three sources (similar results are found for larger communities, see Appendix D), revealing the need to account for these other topological features to better understand the existence of priority effects in ecological communities.

### From theory to testable hypotheses

Our theoretical framework illustrates that priority effects can take a large set of possibilities, most of which are outside the realm of traditional theoretical predictions. However, only empirical evidence can discern which assembly dynamics are possible and which are not, given the internal and external constraints acting on ecological communities (Medeiros *et al.*, 2021). For example, Drake (1991) recorded all the necessary data to empirically map the assembly graph for 3 species of algae (see also Zimmermann *et al.* (2003)), while Warren *et al.* (2003) empirically mapped the assembly graph for 6 species of ciliates. In Figure 5, we analyzed these two assembly graphs following our methodology. The predictions of our framework apply to these two communities. First, they exhibit combinations of dynamical sources (topological features). Second, their predictability increases with more invasions and then saturates to a fixed value. Third, their predictability can be explained from the generic importance (obtained from the trained neural network) of each dynamical source.

Unfortunately, we were unable to locate any other empirical studies that mapped assembly graphs. The dearth of empirical studies regarding these dynamics is not surprising given both the challenges involved in performing detailed experiments and also their under-appreciated potential to answer central questions in community ecology (Fukami, 2015; Vellend, 2016). In this regard, we discuss some viable empirical designs for inferring assembly graphs in ecological communities. A direct approach to infer assembly graphs should include determining which species combinations can persist and then mapping how these combinations change after a new species is introduced. Although this direct approach may seem to require an exhaustive combinatorial design that is too labor-intensive to be feasible, the actual experimental workload can be much lower.

To explain the work required, we first focus on identifying the persistent species combinations (i.e., knowing the nodes in assembly graphs). While a community with *S* species has 2^*S*^ − 1 potential species combinations, it has been shown that only a small fraction of such combinations can persist (Angulo *et al.*, 2020). Because of the sparsity of these persistent combinations, recently developed computational tools based on Bayesian inference (Maynard *et al.*, 2020) and deep learning (Michel-Mata *et al.*, 2021) can facilitate the inference of all persistent combinations using a small number of experiments in a community with *S* species. In brief, the method developed by *Maynard et al.* (2020) can infer the coexistence of all 2^*S*^ − 1 species combinations from a minimum of *S* + 1 experiments based on measures of species’ *absolute* abundances (1 experiment is to grow the full community of *S* species and the other *S* experiments are to grow leave-one-out communities comprising *S* − 1 species each). The goal of the method proposed by Michel-Mata *et al.* (2021) is similar to *Maynard et al.* (2020)’s, but a major difference is that Michel-Mata *et al.* (2021) only requires experimental measures of species’ *relative* abundances. Moreover, if we are only interested in bottom-up assembly (i.e., starting with no species, and then adding species one by one), then persistent combinations that are unreachable via introduction of single species do not need to be mapped to construct the corresponding assembly graph. In Appendix E, we analyzed all assembly dynamics for 3 species and found that more than 90% of assembly dynamics contained unreachable combinations (Figure S7), which would make the inference problem more feasible.

Focusing now on how species composition changes after the introduction of a new species (i.e., knowing the edges in assembly graphs), we suggest that the experimental procedure can follow the standard procedures in assessing the effects of species introductions (Friedman *et al.*, 2017; Grainger *et al.*, 2019a; Spaak & De Laender, 2020; Meroz *et al.*, 2021). In brief, one would introduce an invader species to the resident community at low abundance (relative to the abundance of the resident species) and assess whether species composition changes. Recently developed computational tools can also facilitate the inference (Deng *et al.*, 2021; Pande *et al.*, 2021). If we are only interested in bottom-up assembly, then many edges do not need to be mapped to construct the corresponding assembly graph. In Appendix E, we analyzed all assembly dynamics for 3 species and found that more than 80% of dynamics contained unreachable edges (Figure S8).

While alternative stable states have been the most studied consequence of priority effects, our framework indicates that the three other consequences of priority effects can have a stronger contribution to community predictability, especially in small communities. This possibility can be tested by exploiting the strong constraints between dynamical sources and predictability of priority effects. Specifically, to study these additional dynamical sources, we can use alternative computational approaches based on pairwise interaction strengths inferred from experiments on 2-species communities (Case, 2000). Such empirical data are increasingly available spanning a wide range of study systems, such as annual plants (Godoy *et al.*, 2014; Kraft *et al.*, 2015), perennial plants (Uricchio *et al.*, 2019; Song *et al.*, 2021), and microbial systems (Xiao *et al.*, 2017; Kehe *et al.*, 2020). Thus, the empirical assembly graphs can be computationally mapped with empirically parameterized population dynamics models. However, it is worth remembering that a defining feature of priority effects is that interaction strengths likely depend on assembly history (Fukami, 2015; Song *et al.*, 2018), questioning the validity of inferences based on fixed interaction strengths. Because we know little about how variable these interaction strengths are (Park, 1954), it would be best to combine experimental and computational approaches. That is, the differences between the observed assembly graph in the experimental approach and the inferred graph from the computational approach may provide clues as to how variable interaction strengths are due to the assembly history. A caveat, though, is that these assembly dynamics may take many generations to emerge, thus computational models with fitted empirical interactions from only a few generations risk finding spurious dynamics that do not exist.

## Discussion

In the story “The Garden of Forking Paths,” Jorge Luis Borges envisioned a labyrinth where divergence takes place in time rather than space, and where different paths sometimes lead to the same conclusion. Similarly, an assembly history on an assembly graph can be thought of as a forking path since it also creates temporal trajectories of species composition that can lead to the same final state or different ones. By borrowing tools from graph theory, we have provided a non-parametric framework to scan the complete terrain of the labyrinth of priority effects. This framework has allowed us to classify priority effects, enumerate all possible assembly dynamics operating in a community, and quantify how predictable these dynamics would be if species arrival history was stochastic and unknown.

We have introduced the concept of topologically unique assembly graphs. This concept has allowed us to rigorously estimate the diversity of priority effects in multispecies communities. We have estimated the exact diversity for 2-species and 3-species sets, and, as a first-order approximation, for sets of more than three species (Figure 1D). We have revealed a much richer set of assembly dynamics than traditional parametric approaches typically capture. The parametric approaches typically make two pivotal assumptions: history-independent interaction strength and invasion analysis. In invasion analysis (Grainger *et al.*, 2019b), the invasion criteria assumes only two possibilities for a community with *n* species after invasion: either it has *n* + 1 species (if the invasion was successful), or remains with *n* species (if the invasion was unsuccessful). However, evidence indicates that the set of potential assembly dynamics in ecological communities can be larger than those considered by invasion analyses (Warren *et al.*, 2003; Saavedra *et al.*, 2017; Barabás *et al.*, 2018; Carlström *et al.*, 2019; Amor *et al.*, 2020; Angulo *et al.*, 2020; Deng *et al.*, 2021). Thus, our graph-based approach may provide a more realistic analysis of ecological dynamics than those approaches focusing on history-independent interaction strength and invasion analysis.

Following previous work (Fukami, 2015), we have shown that priority effects can differ in terms of their contribution to community predictability. We have focused this predictability on species composition given the uncertainty derived from the potential assembly histories (Figure 2). We have demonstrated that two types of predictability can be investigated: on the final composition and on the temporary changes in composition (trajectories). We found that these two types are often correlated and yield similar results (Figure 3A). Thus, the two types of predictability have similar information, a phenomena that is analogous to ergodicity in statistics (Strogatz, 2014). However, assembly dynamics is in general not ergodic (e.g., a stable state would render the dynamics non-ergodic) and more research is necessary.

Additionally, we have shown that the predictability of a community generally increases with repeated invasions (Figure 3B). On a conceptual level, with more species invasions, deterministic ecological processes are more likely to overwhelm stochastic events. In the context of assembly graphs, the topological features of assembly graphs become apparent only when when the community trajectories are long enough. Thus, laboratory experiments that only allow single invasions (e.g., Drake (1991); Lawler & Morin (1993); McGrady-Steed *et al.* (1997)), as opposed to multiple invasions (e.g., Robinson & Dickerson (1987)), may underestimate community predictability in nature, where recurrent migrations are common (Newton, 2010; Secor, 2015). The predictability of a community does not always increase with more invasions at the same rate, but rather saturates at a constant value (Figure 3B). Thus, laboratory studies can approximate community predictability in nature with two complimentary approaches. The first approach is to perform a few invasions. About 10 times is more than enough for up to five species (Appendix D), provided that the time interval between invasions is large enough to allow the communities to approach an equilibrium. The second approach is to empirically map the assembly graph, which can fully determine the community predictability with an arbitrarily given pool of assembly history, although it generally takes more experimental efforts.

Moreover, we have shown that this predictability can be used to classify priority effects. We have shown that four dynamical sources can be expressed as topological features within our graph-based approach to know the predictability of assembly dynamics (Figure 4). These sources are the number of alternative stable states, the number of alternative transient paths, the length of compositional cycles, and the presence or absence of an escape from compositional cycles to stable states. We have found that both the frequency and combination of the four topological features are good predictors of how many outcomes to expect (Figure 5). We have also found that the explanatory power of these four topological features increases when species attempt to invade multiple times (Figure 5C). While the number of alternative stable states has received most of the attention (Schröder *et al.*, 2005; Serván & Allesina, 2020; Amor *et al.*, 2020; Abreu *et al.*, 2020), we show that the other three sources, especially in small communities, can contribute more to community predictability (Figure 5D). The trained neural network has predicted the observed predictability of two empirical assembly dynamics (Drake, 1991; Warren *et al.*, 2003) well (Figure 6).

**Figure 6:**
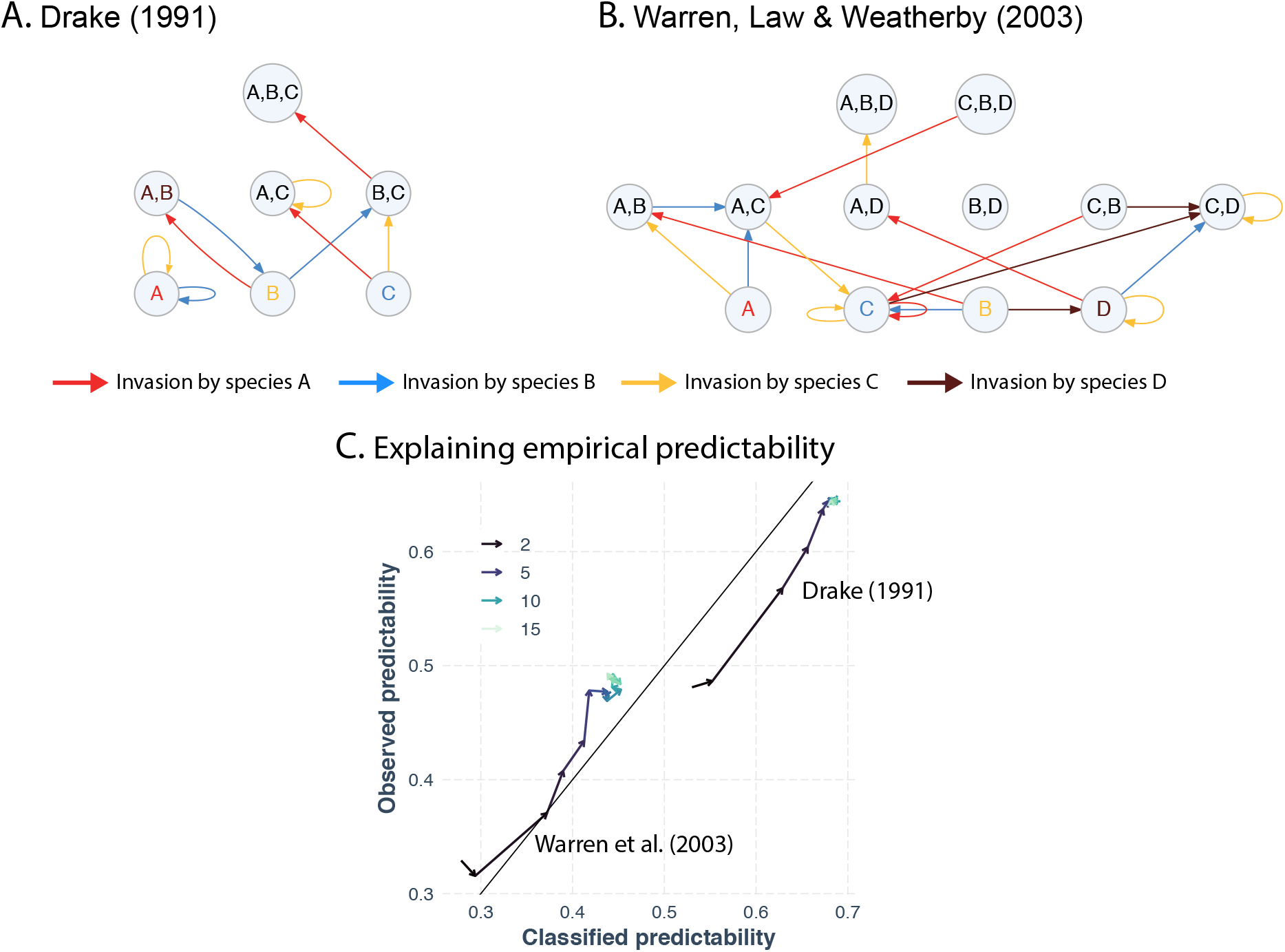
Applying the theoretical framework to empirical data. Panel (**A**) shows the empirical assembly graph for 3 algal species community studied by Drake (1991). Species *A*, *B*, and *C* are *Scenedesmus quadricauda*, *Selenastrum bibrium*, and *Ankistrodesmus falcatus*, respectively. This assembly graph has 3 stable states, 1 transient path, 1 compositional cycle with length 2, and from which the community can escape to a stable state. Panel (B) shows the empirical assembly graph for 4 protist species studied by Warren *et al.* (2003). Species *A*, *B*, *C*, and *D* are *Blepharisma japonicum*, *Colpidium striatum*, *Paramecium caudatum*, and *Tetrahymena pyriformis*, respectively. This empirical assembly graph has 2 stable states, 1 transient path, and no compositional cycles. Panel (**C**) shows the application of our framework on these two empirical examples. Similar to Figure 5A-B, the *x* axis represents the classified predictability (from the test set) using the four topological features, while the *y* axis represents the observed predictability. The color represents the number of repeated invasions (ranging from each species invades 2 times to 15 times). As the the number of repeated invasions increases (the direction of the arrows), the observed predictability of the two communities increase and saturates to a fixed value. The gray line is the 45-degree line, representing that the observed predictability equals the classified predictability. The classified predictability is close to the observed predictability.

Our non-parametric graph-based approach is not intended to replace the parametric model-based approach. Parametric models are irreplaceable tools to understand priority effects, but non-parametric approaches are more flexible for accommodating different theoretical tools (Barabás *et al.*, 2018; Arnoldi *et al.*, 2019; Pande *et al.*, 2020; Spaak & De Laender, 2020). For example, while our approach uncovers three other types of priority effects for 2 species (Figure 1) that are not covered in classic Lotka-Volterra approaches (Fukami *et al.*, 2016; Ke & Letten, 2018), all priority effects can occur in parametric models by integrating processes where the assembly history affects parameters. Thus, our non-parametric graph-based approach can serve as a roadmap for the parametric approach by motivating new modelling strategies of priority effects. A major limitation of our approach is that we have focused on species richness or species composition. The general concept of priority effects also covers the functional properties of ecological communities, such as energy flow and productivity (Fukami & Morin, 2003; Dickie *et al.*, 2012; Tan *et al.*, 2012). A possible solution is to establish a functional map from the nodes or the links in the assembly graph onto the functional property (e.g., productivity function in *Rohr et al.* 2016). The flexibility of our framework may serve as a common currency to characterize priority effects across study systems and theoretical models.

## Acknowledgement

We thank Lucas P. Medeiros, Hengxing Zou, and Pengjuan Zu for insightful discussions. The authors also thank the editor and three reviewers for their suggestions that improved our paper. Funding to S.S. was provided by NSF grant No. DEB-2024349. Funding to T.F. was provided by NSF grant No. DEB-1737758.

